# Replacing reprogramming factors with antibodies selected from autocrine antibody libraries

**DOI:** 10.1101/156349

**Authors:** Joel W. Blanchard, Jia Xie, Nadja El-Mecharrafie, Simon Gross, Sohyon Lee, Richard A. Lerner, Kristin K. Baldwin

**Affiliations:** Department of Molecular and Cellular Neuroscience, Dorris Neuroscience Center, The Scripps Research Institute, 10550 North Torrey Pines Road, La Jolla California 92037, USA; Department of Cell and Molecular Biology, The Scripps Research Institute, 10550 North Torrey Pines Road, La Jolla California 92037, USA

## Abstract

Signaling pathways initiated at the membrane establish and maintain cell fate during development and can be harnessed in the nucleus to generate induced pluripotent stem cells (iPSCs) from differentiated cells. Yet, the impact of extracellular signaling on reprogramming to pluripotency has not been systematically addressed. Here, we screen a lentiviral library encoding ˜100 million secreted and membrane-bound antibodies and identify multiple antibodies that can replace *Sox2/c-Myc* or *Oct4* during reprogramming. We show that one Sox2-replacing antibody initiates reprogramming by antagonizing the membrane-associated protein Basp1, thereby inducing nuclear factors *WT1* and *Esrrb*/*Lin28* independent of *Sox2*. By successively manipulating this pathway we identify three new methods to generate iPSCs. This study expands current knowledge of reprogramming methods and mechanisms and establishes unbiased selection from autocrine antibody libraries as a powerful orthogonal platform to discover new biologics and pathways regulating pluripotency and cell fate.

## Introduction

During development, cellular diversity arises through a series of choreographed cell fate changes that are temporally ordered, largely irreversible and conserved among species. These changes are guided by signals that transit the cell membrane and converge in the nucleus where they rewire transcriptional programs to produce coordinated patterns of gene expression that specify a new cellular identity. Surprisingly, in 2006, Yamanaka and Takahashi found that the transient expression of only a few transcription factors was sufficient to reverse cellular differentiation and produce induced pluripotent stem cells (iPSCs) that recapture the developmental potency of embryonic cells. ^1-4^ Since this discovery, direct reprogramming using transcription factors has emerged as a robust general method to induce cell fate changes that do not normally occur during development such as the direct conversion of fibroblasts into neurons or cardiomyocytes^5-8^. This increased understanding of how to program cell fate from within the nucleus has led to a marked shift in approaches to model or treat disease using human cell types produced *in vitro,* and also informed thinking about how nuclear events may contribute to de-differentiation in other contexts.^9^ In contrast, the potential for membrane-to-nucleus signaling pathways to replace transcription factors during reprogramming has not been systematically addressed.

One barrier to understanding mechanisms underlying induced pluripotency is that reprogramming inefficient and involves stochastic steps, making it difficult to identify pathways involved in successful reprogramming using bulk biochemical analyses or proteomics.^10-12^ Although small molecule and shRNA/CRISPR based screens have uncovered several signaling pathways involved in reprogramming, these screens are limited by their reduced capacity to perturb key signaling modalities such as protein-protein interactions.^13-17^ Furthermore, small molecule based reprogramming has not been widely adopted, perhaps due to the potential for unknown off target effects.^18, 19^ Thus, it is likely that some, or perhaps even the majority of biochemical events that can trigger reprogramming to pluripotency, remain unknown.

One powerful means to establish the function of cell signaling cascades is to employ antibodies to perturb protein function or protein-protein interactions. Historically, synthetic combinatorial antibody libraries have been useful to identify antibodies directed at known targets.^20-28^ Here, however, we wished to identify unknown pathways. To accomplish this we took advantage of the lentiviral gene delivery system. By encoding a synthetic combinatorial antibody library in lentivuses, we can transduce fibroblasts with the library such that each cell will express one or several unique antibody proteins, effectively converting each cell into an autonomous biochemical reaction compartment. Clonal colonies that emerge will contain the gene sequence of a candidate autocrine reprogramming antibody. This sequence can be recovered and used to produce the antibody in soluble, secreted or membrane-bound versions for confirmation of activity and further mechanistic investigations. Therefore, for this study we adapted a previously described combinatorial antibody library for use in unbiased cell based screening with clonal selection of proliferating iPSC colonies. To accomplish this, we generated a lentiviral library in which each lentivirus encodes a unique antibody sequence that is either targeted for secretion or membrane-tethered, with the total library complexity comprising ˜100 million unique specificities.^27, 29-31^ This autocrine antibody platform enables us to survey a very large number of potential independent specificities, which are equal to the number of original cells plated multiplied by the MOI, in this case > 10^8^ specificities in one experiment, without prior knowledge of a target. This platform allows the screening of a large number of molecules compared with previous studies that involved pre-selection of antibodies against a desired target.^32^

Using the autocrine antibody reprogramming platform, we isolated multiple antibodies that replaced either *Sox2/c-Myc* or *Oct4* in generating iPSCs. Identifying the target of one *Sox2* replacement antibody showed that it binds to Basp1, a protein not previously implicated in pluripotency or identified in shRNA screens.^16, 17^ Our mechanistic studies show that Basp1 inhibition leads to increased nuclear *Wt1* activity and increased expression of *Esrrb* and *Lin28*, prior to *Sox2* induction, which differs from previously reported reprogramming pathways, validating the autocrine antibody reprogramming platform as a discovery tool to identify new biologics and pathways that induce pluripotency or impact cell fate plasticity.

## Results

### The autocrine antibody library reprogramming platform

To allow us to select for and recover antibody sequences that promoted clonal growth of pluripotent stem cells from individual transduced fibroblasts without prior selection on a known target, we took advantage of the ability of lentiviruses to insert in the genome of cells. To facilitate lentiviral delivery and sequence recovery, the antibodies were encoded in a single chain form in which the entire recognition domain of the CDR3 region is encoded by a ScFv domain fused to the Fc region from human IgG1, as described previously.^31^ If cells transform into iPSCs they will grow up as clones of single cells that will contain one or more lentiviral insertions **(Figure 1a**). Because the antibody coding sequences are integrated in the genome, their sequences can be recovered by PCR and re-tested for efficacy **(Figure 1a)**. Candidate antibodies can also be targeted for secretion or to various cellular compartments, which can increase their effective concentration. For this study, we a library comprising both secreted and membrane-tethered antibodies **(Figure S1a)**.

**Fig. 1.**
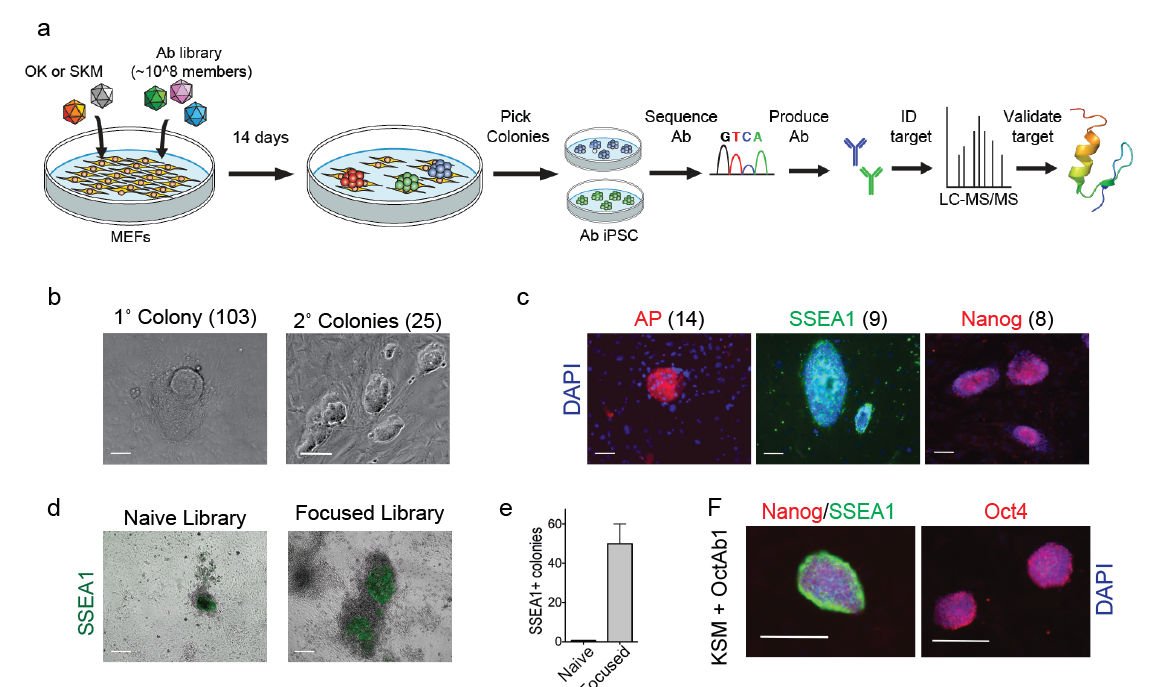
Autocrine selection from combinatorial libraries captures antibodies with reprogramming activity. **a)** Schematic of the combinatorial antibody screening platform **b and c)** Summary and triage of primary colonies from *Sox2* replacement screen. Primary hits were categorized based on their ability to reform colonies following disassociation and the presence of alkaline phosphatase activity, SSEA1, and Nanog expression. **d)** Example of SSEA1-positive primary and secondary colonies observed and captured during the combinatorial screen. Scale bar = 100 µm. **e)** Colonies that contain Oct4-replacing antibodies were selected using the cell surface antigen SSEA1, an intermediate marker of reprogramming. Screening a secondary, Oct4-focused library results in a greater than 100-fold increase in efficiency in generating Oct4 positive colonies (n=3 wells per experiment, p<0.05 t-test, error bars are SEM). **f)** Colonies extracted from Oct4-focused library express pluripotency marker SSEA1, Nanog, and Oct4.

### Identifying antibodies that replace *Sox2*/*cMyc* in reprogramming

We wished to determine whether the autocrine antibody platform could identify antibodies that induce reprogramming without use of the transcription factors *Sox2* and *c-Myc.* To achieve this, we transduced 10^8^ mouse embryonic fibroblasts (MEFs) with doxycycline-inducible lentiviruses encoding *Klf4* and *Oct3/4*, (KO) but not *Sox2* or *cMyc* (SM) and the autocrine antibody lentiviral library (MOI = 0.8) **(Figure 1a)**. This resulted in a population of ˜1 x 10^8^ MEFs, each theoretically expressing one or more unique antibody sequences. In the primary screen, we also included an shRNA targeted against *p53* to increase reprogramming efficiency. This shRNA was not present in the subsequent confirmation experiments. Fourteen days after induction, MEFs infected with the antibody library and the KO transcription factors produced approximately 10 iPSC-like colonies per one million MEFs plated **(Figure S1b)**. We compared these cells to positive control cells expressing all four factors (*Oct3/4, Sox2 Klf4*, and *cMyc,* OSKM) and negative control MEFs transduced with membrane-targeted tdTomato plus KO and *p53* shRNA. Positive controls produced ˜500 colonies per 1 million cells plated, while no colonies arose in the negative control wells **(Figure S1b)**. Because only one copy of each antibody is present, the probability of an antibody with reprogramming activity resulting in detectable pluripotency may be limited by the low efficiency of reprogramming (0.1 - 2%). Therefore, we initially screened using a low stringency metric (colony formation) to capture 103 iPSC-like colonies and expanded them in independent wells **(Figure 1b)**. After single cell dissociation, twenty-five of the primary colonies gave rise to secondary colonies that could be propagated as cell lines under standard ES/iPS cell culture conditions. Next, to identify bona fide potential iPSC colonies tested them for increasingly stringent pluripotency markers, alkaline phosphatase, SSEA1 and Nanog and identified eight triple positive colonies **(Figure 1c)**. Notably, more colonies expressed the early (AP) and intermediate (SSEA1) markers than Nanog, while all Nanog positive colonies were triple positive, suggesting that antibody induced reprogramming was following previously reported steps.

### Two-step autocrine antibody selection identifies antibodies that replace *Oct4*

We next examined whether the autocrine antibody platform could identify antibodies that replace *Oct4* in reprogramming, which has proven challenging in other studies.^33, 34^ Therefore, to increase our chance of success, we performed a two-step screen with a design parallel to the *Sox2/c-Myc* screen, except that cells were transduced with *Klf4*, *Sox2*, and *cMyc* (KSM) but not *Oct4* **(Figure 1a)**. Since the Sox2 screen suggested that SSEA-1 is a representative early indicator of reprogramming we selected the ˜50 SSEA positive colonies we found in a screen of ˜10^8^ MEFs (no positives were present in KSM only wells) **(Figure. 1d and e)**. We next used PCR to amplify these antibody sequences and subcloned them into new lentiviruses to generate a secondary *Oct4*-focused library **(Figure 1a)**. We next repeated the original screen using the focused library, which generated more than 50 SSEA1-positive colonies/10^6^ cells, representing a ˜100-fold increase in activity over the naïve library **(Figure 1e)**. We picked individual colonies from the focused screen and clonally expanded them into lines. Cell lines established from the focused library expressed the pluripotency markers Nanog and Oct4 **(Figure 1f)**. These experiments validate two different strategies for employing the AutoAb platform to identify antibodies that replace reprogramming factors.

### Validation of iPSCs generated using antibodies to replace Sox2 or Oct4

To functionally validate the antibody hits, we used genomic PCR to recover ScFv antibody coding sequences from two independent Nanog-positive secondary lines from each screen. DNA sequencing showed that each line harbored a unique CDR3 region **(Figure S2a and b, Table S1)**. To confirm their function, we subcloned these sequences into two different viral backbones, containing either signal sequences for secretion or for membrane tethering (MTA). These lentiviruses were then tested in new reprogramming assays. When targeted to the membrane, both *Sox2* replacing antibodies (SoxAb1 and SoxAb2) and *Oct4* replacing antibodies (OctAb1 and OctAb2) produced Nanog-positive colonies in the absence of *Sox2* or *Oct4* **(Figure 2a and b).** Similar results were observed with lentivirally-encoded secreted forms of the antibodies (data not shown).

**Fig. 2.**
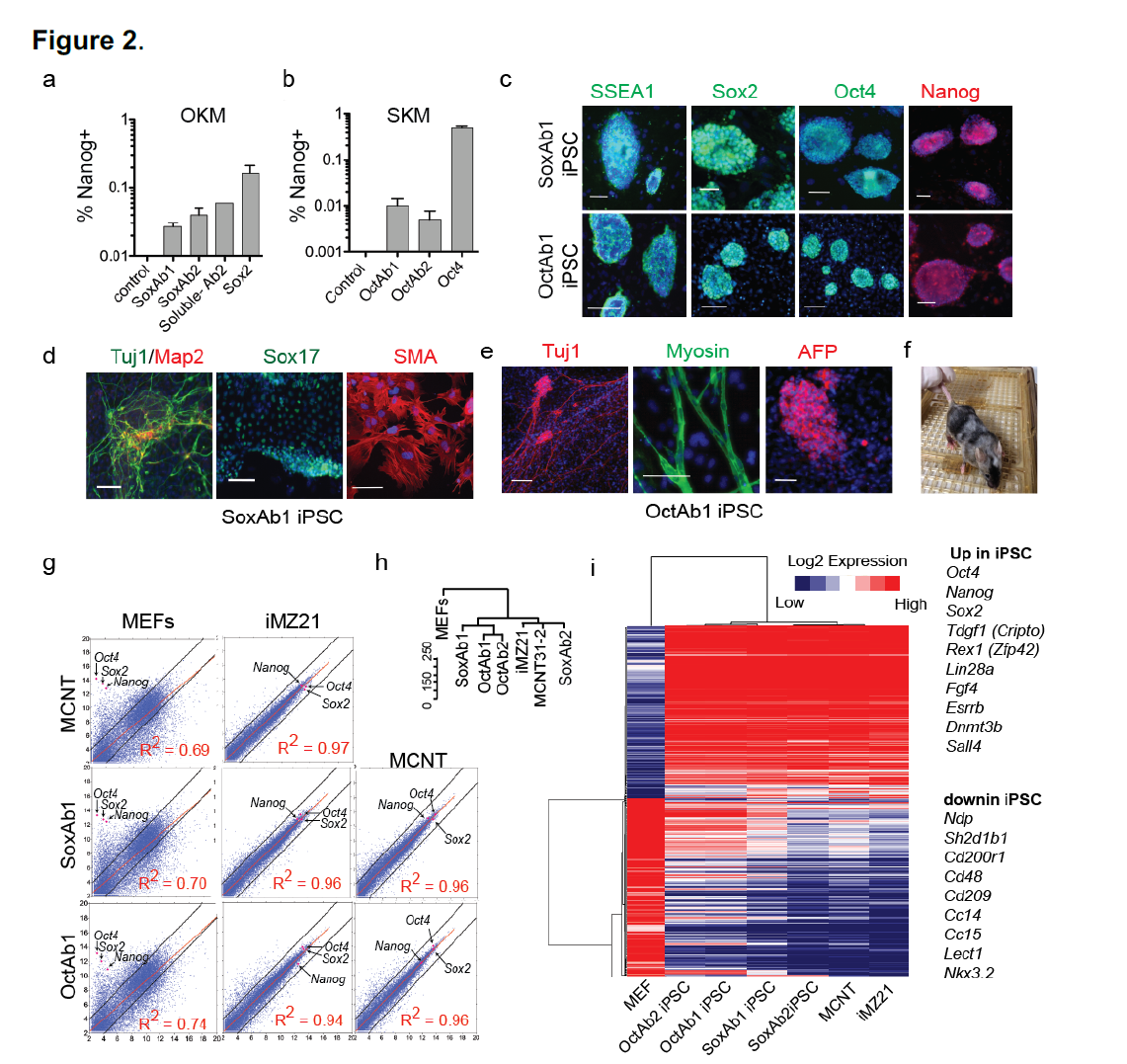
Antibodies enable reprogramming without Sox2 and Oct4. **a)** Membrane-tethered antibody (MTA) clones 14 and 39 (SoxAb1 and SoxAb2) and soluble SoxAb2 enable reprogramming in the absence of Sox2. Bars represent means from three biological replicates, except for the soluble Sox2 antibody (n=1 experiment due to limiting antibody). Error bars = SEM. **b)** MTA8 and MTA20 (OctAb1 and OctAb2) reprogram fibroblasts in the absence of *Oct4*. Bars represent means from three biological replicates. Error bars = SEM. **c)** iPSC lines generated with SoxAB1 + Klf4 and Oct4 (minus Sox2 and cMyc) and OctAb1 + Klf4, Sox2, and cMyc (minus Oct4) appropriately express pluripotency markers SSEA1, Sox2, Oct3/4, and Nanog. Scale bars = 100 µm. **d and e)** iPSC lines generated with SoxAb1 and OctAb1 differentiate *in vitro* to the three embryonic germ layers. Scale bars = 100 µm. **f)** iPSC lines generated with SoxAb1 or SoxAb2 differentiate *in vivo* and contribute to mouse tissues as shown by the picture of an eight-week-old chimeric mouse formed by injection of KO + SoxAb1 cells (CD1 genetic background) into a C57BL6 blastocyst. **g)** RNAseq scatter plots showing global gene expression of antibody iPSC lines is highly similar to TEC competent iPSC (iMZ21) and mESC (MCNT) lines but very different from somatic MEFs. **h)** Dendrogram tree depicting transcriptional similarity based on whole genome hierarchical clustering (uncentered correlation with complete linkage). **i)** Heatmap depicting 400 differentially expressed genes between MEFs and TEC-competent iPSC (iMZ21) and mESC (MCNT) determined by RNA-sequencing. Each column represents mean of three biological replicates for each line. Red and blue indicate high and low levels of expression respectively. Genes and samples are organized by hierarchical clustering of 400 differentially expressed genes (using uncentered correlation with complete linkage).

While lentiviral encoded antibodies are facile tools to identify targets and pathways relevant for reprogramming, we also wished to test whether exogenous application of soluble versions of the antibodies identified in this screen could induce reprogramming. To test this we produced a soluble form of SoxAb2 antibody and showed that adding this protein directly to the culture media (at ˜1.0 ug/ml) can also induce reprogramming in the absence of *Sox2* **(Figure 2a)**. This serves as a proof of concept that antibodies identified using lentivirally encoded secreted molecules and also have activity when produced exogenously. However, as these initial antibody hits are likely to serve as a starting point for optimizing exogenously applied antibodies that would need to be optimized for protein production, biding affinity, etc. we did not further address this mechanistically. Finally, we also tested the lentiviral Sox2 or Oct4 replacing antibodies with transcription factor cocktails lacking Oct4 or Sox2 respectively, and observed no colonies (data not shown), indicating that the antibodies we identified exhibit pathway based selectivity.

To demonstrate conclusively that antibodies can replace *Sox2/c-Myc* or *Oct4* in reprogramming to pluripotency we performed a series of functional tests of the iPSC lines **(Table 1 and Table S1)**. iPSCs generated with the subcloned and sequenced *Sox2* or *Oct4* replacing antibodies could be expanded into cell lines that maintained morphological and self-renewal properties similar to mouse ESCs and four factor iPSCs **(Figure S2c and d)**. PCR with primers specific to viral *Oct3/4*, *Klf4*, *Sox2,* and cMyc, confirmed that iPS cell lines generated with Sox2 antibodies or Oct4 antibodies did not mistakenly harbor transgenic *Sox2/cMyc* or *Oct4* respectively **(Figure S2e and f).** Antibody reprogrammed cell lines all expressed the pluripotency markers alkaline phosphatase, SSEA1, endogenous Sox2, endogenous Oct3/4, and Nanog **(Figure 2c and S2g)**. Antibody iPSC lines also formed embryoid bodies that differentiated into tissue from all three embryonic germ layers including contractile cardiomyocytes (**Figure 2d, 2e, S2h, and Supplemental video 1-3)**.

**Table 1.** Summary of blastocyst injection results

| Line | Blasts transferred | Live pups | Degree of coat color chimerism |  | Germ line transmission |
| --- | --- | --- | --- | --- | --- |
|  |  |  | Low | High |  |
| SoxAb1 | 120 | 9 | 2 | 1 | Not tested |
| SoxAb2 | 100 | 12 | 1 | 1 | Not tested |
| KOMWT1.1 | 111 | 17 | 1 | 3 | + |

We next tested whether these lines could contribute to three germ layers in chimeric mouse embryos. iPSCs derived from albino CD1 fibroblasts using either SoxAb1 or SoxAb2 were injected into C57/BL6 blastocysts (pigmented background). Pups showed significant contribution across all three embryonic germ layers and survived to adulthood **(Figure 2f, S2i, S2j, and Table 1).** These results demonstrate that iPSCs produced with antibodies can pass the standard functional tests used to designate iPSCs as pluripotent stem cells.

To more comprehensively characterize the iPSCs we performed RNA-Seq on triplicate samples from the SoxAb and OctAb iPSC lines and compared them to several gold standard pluripotent cell lines generated in our laboratory that can produce fertile adult mice in tetraploid embryo complementation (TEC) assays (one iPSC line produced with using OSKM, the other produced from a neuron via cloning by somatic cell nuclear transfer (SCNT)). The global transcriptional profiles of the candidate iPSCs that were reprogrammed with antibodies were as similar to TEC competent mouse iPSC (iMZ21) and SCNT-ESC lines (MCNT) as the TEC lines were to each other, and all were similarly divergent from MEFs (Pearson correlation coefficient = 0.94 - 0.96) (**Figure 2g, 2h, S2l).**^1^ Cluster analyses of the global transcriptional profiles suggested that the line produced with SoxAb2 bears the strongest similarity to the TEC competent iPSCs, although this was a subtle difference **(Figure 2h).** To identify any potentially problematic transcriptional changes we used DeSeq to identify a set of key IPSC genes (the 400 genes most differentially expressed between MEFs and the TEC competent lines). Generating a heat map for all lines shows that the “core” iPSC genes that are most upregulated in high-quality lines are also similarly upregulated in the SoxAb and OctAb lines, with some limited residual MEF gene expression detectable in a few of the Ab lines **(Figure 2i).** We also identified genes differentially expressed between each SoxAb or OctAb line and the TEC competent PSC lines, which identified very few (˜100-300) differentially expressed genes **(Figure S2m).** Inspection of these gene lists did not uncover dysregulation of any known pluripotency genes **(Table S2).** Quantitative RT-PCR experiments also confirmed that pluripotency genes (*Oct4*, *Sox2*, *Nanog*, *Lin28*, *Sall4*, *Esrrb*, *Dppa2*, and *Zscan4b*) were upregulated in SoxAb1 and 2 iPSC lines relative to fibroblasts expressing KOM alone **(Figure S2k).** These results show that multiple antibodies selected from combinatorial libraries can replace transcription factors in producing iPSC lines that exhibiting the functional and molecular hallmarks of high-quality pluripotent stem cell lines.

### SoxAb2 binds to and antagonizes Basp1

Next, we wished to determine whether we could use these antibodies to identify new pathways that promote reprogramming to pluripotency. The SoxAb2 iPSCs were the most similar to the TEC competent PSCs and we were able to purify sufficient amounts of this antibody to show that it can act as a biologic when supplied in the reprogramming dish (Figure 2a), making it an attractive candidate to follow up mechanistically. However, Sox2 has previously been replaced by inhibiting Tgf-β signaling, inhibiting Src-kinase or antagonizing the induction of mesoderm genes.^13, 15, 35, 36^ To determine whether the SoxAb2 involved these known mechanisms, we performed a series of experiments that showed that SoxAb2 did not impact Tgf-β signaling (Smad2/3 phosphorylation), Src-kinase activity, or antagonize the induction of mesoderm genes (*Eomes*, *T*, and *Ctnnb1*) **(Figure 3a, Figure S3a, S3b, and S3c)**. These data predict that identifying the target of the SoxAb2 could uncover a previously unrecognized pathway to pluripotency.

**Fig. 3.**
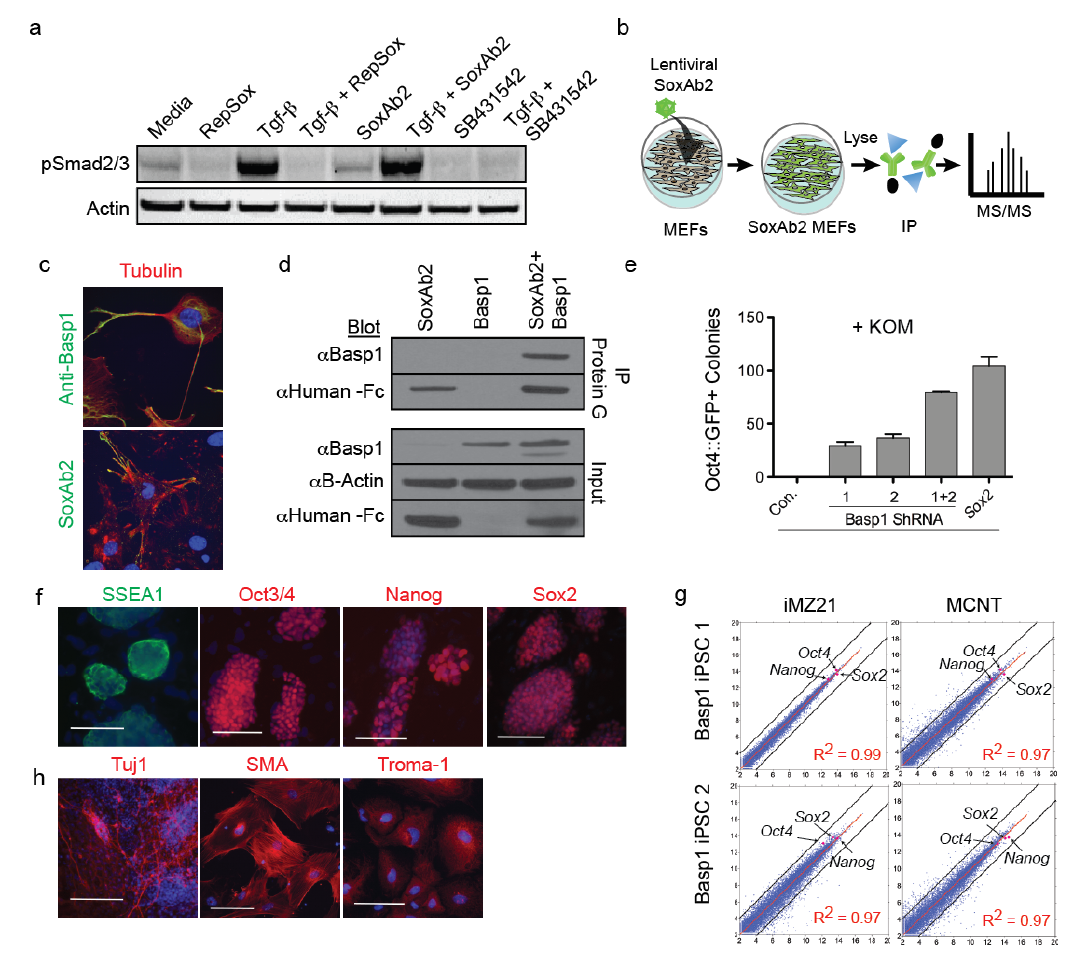
SoxAb2 antagonizes Basp1. **a)** SoxAb2 does not inhibit Tgf-β induced phosphorylation of Smad2/3. In contrast, small molecule inhibitors RepSox and SB431542 previously reported to promote Sox2-independent reprograming inhibit Tgf-β induced phosphorylation of Smad2/3. **b)** Schematic depicting strategy for target identification. **c)** SoxAb2 and Basp1 exhibit similar sub-cellular localization in MEFs. Basp1/SoxAb2 antibody staining is shown in red, tubulin staining in green, and nuclei are stained with DAPI (blue). **d)** SoxAb2 immunoprecipitates Basp1. HEK cells were transduced with SoxAb2 (lanes 1,3) or/and Basp1 (2 and 3 respectively). Lysates were immunoprecipitated with protein G and Western blots were performed with the specified antibodies. **e)** shRNA-mediated knockdown of *Basp1* permits reprogramming in the absence of *Sox2*. Basp1 shRNA 1 and 2 are two independent shRNAs targeted against Basp1. Bars are means from three biological replicates. Y axis represents Oct4::GFP positive clones per 50,000 cells. Error bars are SEM. All treated samples are significantly different from the control by ANOVA followed by Newman-Keuls post-hoc analysis. **f)** *Basp1* knockdown iPSC lines express pluripotency markers SSEA1 (green), Oct3/4, Nanog, and Sox2 (all in red). Blue is nuclear staining with DAPI. Scale bars = 100 µm. **g)** RNAseq scatter plots showing similarity of global gene expression of Basp1 Knockdown iPSC lines compared to TEC competent iPSC (iMZ21) and mESC (MCNT) lines. **h)** *Basp1* knockdown iPSCs differentiate in vitro to the three embryonic germ-layers as shown with antibodies against Tuj1 (neuronal ectoderm), SMA(mesoderm) and Troma-1 (endoderm). Antibody staining is presented in red, blue is nuclear staining with DAPI. Scale bars = 100 µm.

Therefore, we over-expressed SoxAb2 in MEFs and immunoprecipitated with protein G **(Figure 3b)**. SDS/PAGE revealed a unique protein band of ˜25 kDa in MEF lysate expressing SoxAb2 when compared to negative control MEFs lysates expressing *cMyc* or with no infection **(Figure S3d)**. Mass spectrometry analysis of trypsin digests of this band identified several proteins enriched in this band compared to controls: After vimentin, which is a common contaminant in mass spectrometry, the most abundant protein was brain acid soluble protein 1 (Basp1) **(Figure S3d).**

Basp1 is a 22 kDa calmodulin-binding protein that is localized to lipid insoluble membrane domains and cellular outgrowths such as pseudopodia and filopodia ^37-39^ (Korshunova et al., 2008; Maekawa et al., 1993; Mosevitsky et al., 1997). To test whether SoxAb2 bound to Basp1 we compared their patterns of immunostaining in MEFs and performed biochemical experiments. SoxAb2 immunostaining localized to the membrane and cellular processes of MEFs, similar to Basp1 staining **(Figure 3c)**. In addition, we showed that SoxAb2 immunoprecipitates both native and overexpressed Basp1, indicating that Basp1 is present in MEFs at the membrane and is a direct target of SoxAb2 **(Figure 3d and S3e)**.

Antibodies frequently act as antagonists by blocking protein-protein interactions. However, some antibodies can have agonist or catalytic activities. To test whether SoxAb2 was likely to antagonize Basp1 function we compared its effect on MEF gene expression to that of reducing Basp1 expression using shRNAs. After expressing SoxAb2 or Basp1 shRNA in MEFs for 4 days in the presence or absence of the KOM factors we used RT-PCR to assess the induction of selected reprogramming and pluripotency-associated genes. The transcriptional patterns of MEFs expressing either SoxAb2 or Basp1 shRNAs were highly correlated in the absence of the KOM factors and resulted in the upregulation of genes associated with reprogramming such as *Cdh1* (also known as E-cadherin) **(Figure S3f-h)**.^40, 41^

### Inhibiting Basp1 expression promotes reprogramming

If SoxAb2 promotes reprogramming by antagonizing Basp1 then replacing Sox2 with *Basp1* shRNA should also permit reprogramming with KOM. Indeed, mixtures or independent expression of two different shRNAs targeted against *Basp1* efficiently generated *Oct4*::GFP-positive colonies in the absence of ectopic *Sox2* **(Figure 3e)**. These Oct4::GFP-positive colonies were expanded into lines that propagated for at least ten passages under standard mESC culture conditions while retaining properties of pluripotency **(Figure S3i and Table S1)**. PCR with primers specific for *Klf4*, *Oct4*, and *Sox2* transgenes confirmed that *Basp1* knockdown lines did not harbor transgenic *Sox2* **(Figure S3j)**. Similar to SoxAb2 iPSC lines, *Basp1* knockdown lines expressed the pluripotency markers SSEA1, Oct4, Nanog, and endogenous Sox2 **(Figure 3f)**. The global transcriptome profile of cells reprogrammed with Basp1 knock-down was similar (Pearson correlation coefficient = 0.99 and 0.96) to fully pluripotent iPS and ES cells 1(Boland et al., 2009) **(Figure 3g and S3k).** These iPSCs also formed embryoid bodies that contained differentiated cell-types from the three distinct embryonic germ layers including contractile myocytes **(Figure 3h and Supplemental video 4).** Together, these results demonstrate that inhibition of *Basp1* is sufficient to reprogram cells to a pluripotent state in the absence of ectopic *Sox2* expression.

### Basp1 inhibits reprogramming through repression of Wt1

Knocking down Basp1 and expressing an antibody that binds to Basp1 result in similar cell reprogramming outcomes, suggesting that Basp1 may repress an activator of reprogramming. In some cell types, Basp1 has been shown to translocate from the membrane to the nucleus where it acts as a transcriptional repressor of Wilm’s tumor suppressor 1 (Wt1).^42, 43^ Therefore, we hypothesized that SoxAb2 binding to Basp1 or (reduction of Basp1 via shRNA) de-repress Wt1, leading to Wt1-mediated transcriptional activation and reprogramming.

To test this hypothesis we asked whether Basp1 interacts with Wt1. Immunoprecipitation experiments revealed that Basp1 interacts with Wt1 in MEFs **(Figure S4a)**. Since Wt1 and Basp1 interact in cells undergoing reprogramming, we next tested whether Wt1 expression can also replace Sox2 in reprogramming to pluripotency. Indeed, overexpression of *Wt1* (the KTS-minus isoform known to act as a transcription factor) induced *Oct4*::GFP^+^ colonies in the absence of Sox2 **(Figure 4a)**. Next, we established iPSC lines generated with *Wt1* instead of Sox2 **(Table 1 and S1).** These *Wt1* iPSC lines displayed typical mESC morphology and expressed endogenous pluripotency genes **(Figure 4b, S4b, and S4c)**. The global transcriptional profiles of WT1 iPSCs were highly similar to TEC competent iPSCs and ESCs and distant from MEFs **(Figure 4c, S4d, and S4e)**. Functional tests showed that Wt1 iPSCs can generate cells from all three embryonic germ layers *in vitro* and can also contribute to chimeric mice with germline transmission **(Figure 4d and S4f)**. These results demonstrate that *Wt1* can replace Sox2 in generating functional iPSCs, supporting the idea that Basp1 inhibition acts through de-repression of WT1, either by directly blocking protein protein interactions, inhibiting Basp1 translocation or an alternative mechanism.

**Fig. 4.**
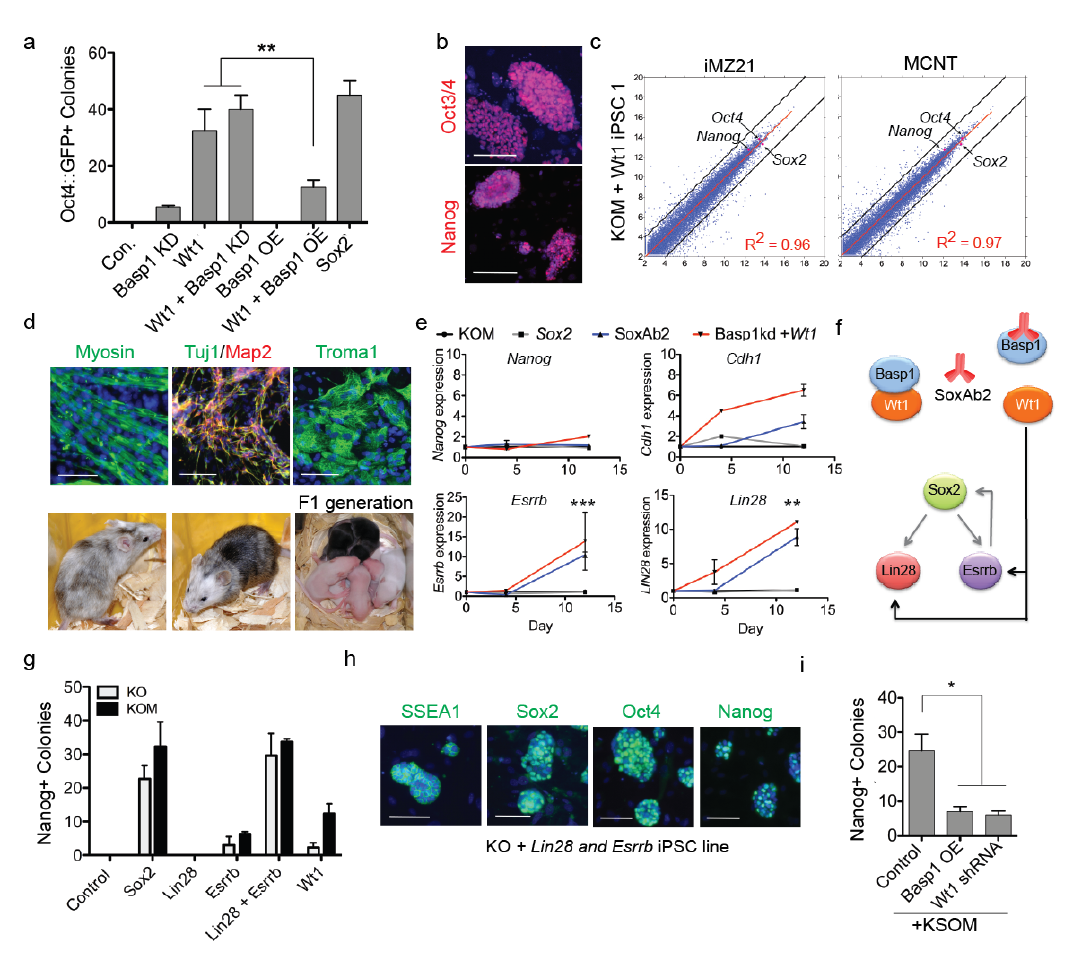
Basp1 repression promotes reprogramming via Wt1-mediated upregulation of Lin28 and Esrrb. **a)** Overexpressing *Wt1* permits reprogramming in the absence of Sox2. *Wt1* reprogramming efficiency is significantly reduced when co-expressed with Basp1. Bars are means from three biological replicates. Error bars are SEM. Significance was tested using ANOVA followed by Newman-Keuls post-hoc analysis, * represents p<0.05. **b)** KOM + Wt1 iPSC lines express pluripotency markers Oct3/4, Nanog, (red). Nuclei are stained with DAPI (Blue). Scale bars = 100 µm. **c)** RNAseq scatter plots showing global gene expression of KOM + Wt1 iPSC lines are highly similar to tetraploid competent iPSC (iMZ21) and mESC (MCNT) lines but very different from somatic MEFs. **d)** KOM + Wt1 iPSC lines differentiate *in vitro* to the three embryonic germ-layers denoted by myosin expression (mesoderm, green), ectodermal markers Tuj1 (green) and Map2 (red), and Troma-1 (green), a marker of endoderm. Nuclei are stained with DAPI (Blue). Scale bars = 100 µm. *In vivo* Wt1 iPS cells contribute to live chimeric offspring with germ-line transmission. CD1 (albino) iPS cells where injected into C57/BL6 (pigmented) blastocysts. White fur indicates the contribution of the iPSC lines. **e)** Expression of *Nanog*, Cdh1, *Essrb* and *Lin28* in KOM-expressing MEFs following continuous treatment with Wt1/Basp1-shRNA, SoxAb2, or Sox2 for four or twelve days. All expression levels are normalized to KOM alone (Control). Bars represent composite means from three biological and four technical replicates. **P < 0.01. *** P < 0.0001. Bonferroni corrected significance level for multiple comparisons q = 0.0166. **f)** Schematic model of an alternative pathway to produce iPSCs. In this SoxAb2 promotes reprogramming by preventing Basp1 from repressing Wt1. Wt1 then activates Lin28 and Esrrb, but not Sox2. Ultimately Sox2 becomes activated by a feed-forward transcriptional network involving Esrrb and stabilizes in a positive feedback regulatory network involving Sox2, Esrrb and Lin28. **g)** Lin28/Esrrb and Esrrb (alone) promote reprogramming in the absence of Sox2. Bars represent means from three biological replicates. Error bars = SEM. **h)** iPSC lines generated with Lin28/Esrrb + Klf4 and Oct4 express pluripotency markers SSEA1, Sox2, Oct4, and Nanog. **i)** Overexpression of Basp1 and Wt1 knockdown significantly impairs OSKM mediated reprogramming into Nanog-positive colonies. Bars represent means of three biological replicates. (significance determined by ANOVA, * p<0.05).

To further test this idea, we asked whether Basp1 and Wt1 exhibit synergistic or antagonistic activities during reprogramming. When *Wt1* and *Basp1* shRNAs were co-expressed, the number of Oct4::GFP^+^ colonies did not significantly increase, suggesting that Basp1 and Wt1 act in the same pathway **(Figure 4a)**. Conversely, when *Basp1* was overexpressed with *Wt1*, the number of Oct4::GFP^+^ colonies was significantly reduced by 68% ± 0.063% (p < 0.0003) demonstrating that Basp1 can interfere with Wt1reprogramming activity **(Figure 4a)**. These results are consistent with a model in which Basp1 in fibroblasts represses Wt1 activity and when this repression is relieved by SoxAb2 or by knocking down Basp1, Wt1 activates transcriptional networks that promote reprogramming.

### Wt1 induces expression of *Lin28* and *Esrrb* during reprogramming

To examine the mechanism by which the Basp1-Wt1 signaling pathway promotes reprogramming we wished to examine early molecular events during the reprogramming process. These can be difficult to detect because of the low efficiency at which cells are reprogrammed.^11, 44^ However, rare events during reprogramming can be detected by quantitative RT-PCR. Therefore, we used RT-PCR to quantify expression of 38 genes associated with reprogramming in KOM MEFS treated with SoxAb2, *Wt1/Basp1*-shRNA or *Sox2* using real-time qRT-PCR. The Basp1/Wt1 pathway might alleviate the need for *Sox2* overexpression by altering cell cycle or by upregulating endogenous *Sox2*. However, Wt1/Basp1 treatment did not alter expression of cell cycle regulators or detectably upregulate endogenous *Sox2* during early or late stages of reprogramming **(Figure S4g)**. Alternatively, Basp1/Wt1 signaling could replace *Sox2* by inducing genes downstream of *Sox2* that ultimately lead to Sox2 induction such as *Nanog*.^13^ However, we did not observe significant upregulation of *Nanog* at day 4 or 12 of reprogramming, further suggesting that the Basp1/Wt1 pathway identified by SoxAb2 operates through a novel mechanism **(Figure 4e)**.

In contrast, *Lin28* and *Esrrb* were significantly upregulated in KOM cells after twelve days of treatment with either SoxAb2 or Wt1/Basp1-shRNA **(Figure 4e)**. *Cdh1* was also upregulated in cells treated with SoxAb2, or Wt1/Basp1-shRNA, but did not reach significance when corrected for multiple comparisons **(Figure 4e)**. Both *Lin28* and *Esrrb* have been previously shown to be downstream of *Sox2* during reprogramming.^45-47^ Furthermore, *Esrrb* has been reported to positively regulate *Sox2* expression, suggesting a possible mechanism by which endogenous *Sox2* is ultimately activated during Basp1/Wt1 reprogramming.^47^

These data support a model in which antibody-mediated antagonism of Basp1 converts Wt1 to a transcriptional activator and leads to expression of *Lin28* and *Esrrb* (either directly or indirectly **(Figure 4f)**. This model predicts that *Lin28*/*Esrrb* and to a lesser extent *Esrrb* alone should promote reprogramming in the absence of *Sox2*. Supporting this model, *Esrrb* alone was able to promote reprogramming with low efficiency and co-expressing *Esrrb* with *Lin28* produced similar numbers of colonies as Sox2 itself **(Figure 4g).** IPSC lines generated with *Lin28*/*Esrrb* or *Esrrb* expressed pluripotency markers and could be propagated under standard mESC culture conditions **(Figure 4h and S4h).** This demonstrates that *Lin28* and *Esrrb* can also promote reprogramming in the absence of *Sox2,* supporting our model.

Inhibiting Basp1 or increasing Wt1 signaling can promote reprogramming in the absence of ectopic *Sox2*. This raises the question of whether Basp1 overexpression or WT1 reduction would inhibit reprogramming. To address this, we derived MEFs from an optimized reprogrammable mouse strain in which KOSM can be uniformly induced using doxycycline.^10^ Overexpression of *Basp1* and knockdown of *Wt1* significantly reduced the number of Nanog-positive colonies by 71.6% ± 0.06% and 75% ± 0.04% (p < 0.0081) respectively **(Figure 4i)**. Similarly, knockdown of *Wt1* reduced *Esrrb*, *Lin28* and its downstream target *cMyc* in mESCs (v6.5) and iPSCs **(Figure S4i).** These experiments show that these pathways intersect and suggest that high expression of Basp1 and/or low Wt1 activity can impede reprogramming by the KOSM factors.

Here we identified a new candidate pathway involving Basp1 inhibition/reduction, Wt1 activation, and Esrrb/Lin28 activation that is involved in reprogramming to pluripotency. WT1 might influence LIN28 gene expression directly or indirectly. Previous ChIP-sequencing studies have determined the consensus DNA binding sequence for WT1 and found that it preferentially binds regions distal to the transcription start site (TSS) with the majority of binding occurring ±50 – 500 Kb (Motamedi et al., 2014). We performed *in silco* analysis of *LIN28A* and *B* genomic region and found the presence of WT1 binding sites at distal regions of each gene **(Figure S5).** Using ChIP-qRT-PCR experiments we show enrichment of WT1 in these regions relative to a non-specific IgG control **(Figure S5).** In contrast, a gene that is not regulated by WT1 (RPL30) and a genomic region void of genes we did not observe enrichment of WT1 over control. These results show that WT1 occupies genomic regions distal to *LIN28A* and B TSS and, therefore, could directly regulate their transcription.

Together, these experiments show that the autocrine antibody platform can identify new pathways relevant to cell fate plasticity that operate in artificial environments such as induced pluripotency as well as in endogenous reprogramming situations such as cancer progression.

## Discussion

This study validates the autocrine antibody platform for use in the discovery of biologics and pathways that induce pluripotency, building on powerful studies of other groups that have used pre-selected antibody libraries to alter ES cell fate.^32^ Together these studies highlight the power of this approach in diverse cell based screening contexts. For example, by performing these screens we identify at least three new methods to produce iPSCs. In addition, we establish that perturbing membrane-to-nucleus signaling pathways is a useful method to identify new mechanisms that induce pluipotency, which would perhaps not have been predicted due to the success and wide adoption of transcription factor-based reprogramming methods acting directly in the nucleus.

Here, we identify antibodies that can replace *Sox2/c-Myc* and *Oct4* during reprogramming, but we have not yet been able to replace *Klf4*. Replacing three of the four reprogramming factors with antibodies suggests that it may one day be possible to induce pluripotency solely through the use of antibodies. Fully antibody-based induction of pluripotency could offer several practical advantages. First, applying soluble antibodies to cells in solution offers a means to ensure that each cell receives the same biochemical trigger as every other cell, which is difficult to accomplish with methods that rely on cell transduction or transformation. Second, using soluble antibodies to reprogram cells could provide a facile method to generate iPSCs that have not experienced any direct nuclear perturbation, thus limiting threats to the iPSC genome. As such, RNA transfection has emerged as a popular non-integrating reprogramming approach. However, RNA-based reprogramming requires frequent transfection of the cells and individual iPSCs are likely to have experienced different ratios and levels of reprogramming factors which could lead to increased variability among iPSC lines.^48^

Other groups have used cocktails of small molecules to induce pluripotency in mouse cells.^33^ While an appealing approach, small molecules can exhibit unknown off-target effects at the concentrations required. In contrast, antibodies typically bind their antigens with high specificity and have been shown to exhibit >200-fold improvements in selectivity when compared to small molecules aimed at the same target.^49^ Finally, once a target is identified, antibodies can be readily “evolved” to produce molecules with increased binding affinity or stability. Together these unique features of antibodies and our demonstration that multiple soluble and membrane-tethered antibodies can induce reprogramming, establishes this as a promising new avenue for research. Although we have not yet identified an antibody to replace *Klf4* in reprogramming, we believe that these studies support the initiation of larger scale screens and additional analyses of reprogramming pathways may ultimately identify a non-integrating all-antibody method for reprogramming. Alternatively, if reprogramming entirely with antibodies is not possible, reprogramming with a mixture of antibodies and other chemical or RNA-based inducers may still result in iPSC production methods with improved safety or reproducibility.

Another advantage of reprogramming with genetically encoded antibodies is that this approach is compatible with several well-validated methods to improve antibody efficacy and thus increase reprogramming efficiency. One method, akin to phenotypic screening, is to mutagenize the identified antibodies through PCR or similar approaches, then perform new selections for iPSCs. This approach can identify antibodies with the most effective balance of kinetic and structural features and thus rapidly optimize reprogramming efficiencies, even without knowledge of the antibody ligand. In cases where the ligand of the antibody is abundant enough for target identification, the power of combinatorial antibody library screening can be further exploited. By screening for antibodies that bind to a known ligand, either via traditional phage display, or through cell-based screening, one can identify new antibodies directed to the same ligand that may have very different binding sites or mechanisms. Importantly, using several non-overlapping antibodies in the same cocktail might lead to further improvements in reprogramming efficiency. It is also possible that the inherent stochastic nature of reprogramming may be due to our lack of knowledge of more efficient pathways than those engaged by OSKM mediated reprogramming. In this case, wider adoption and continued application of the unbiased Autocrine antibody library platform may uncover new reprogramming mechanisms that are inherently more efficient than current protocols.

## Experimental Procedures

### Cell culture and iPS cell generation

All animal research was performed with the oversight of the Office of Animal Resources at The Scripps Research Institute. MEFs were prepared from CD1, or a reprogrammable mouse line (Jackson Laboratories, Stock number 011004) as previously described.^50^ Lentiviruses was produced in HEK-293 cells grown on flasks coated with 0.0001% poly L-lysine in DMEM supplied with 10% FBS. Each flask was transfected with a solution of 850 µl HBSS, 100 µl 2M CaCl_2_, transfection plasmids RRE (15 µg), REV (15 µg), pDMG.2 (15 µg) and lentiviral expression vector (15 µg). The media was replaced after 24 hours with DMEM supplied with 30% FBS and after 48 hours the supernatant collected, spun for 5 minutes at 200 g and filtered through 0.45 µm PVDF filters. Lentiviral infections were performed as previously described.^50^ Expression was induced with 5 µg/ml doxycycline. The first day of doxycycline addition was denoted as day 0 of expression. For the first seven days cells were cultured in DMEM supplemented with 10% fetal bovine serum (FBS). After 7 days cells were cultured in mES cell media supplemented 15% Knock-out serum replacement, LIF. All colony counts were performed on day 14 unless noted otherwise.

### Gene knockdown and overexpression validation

ShRNAs were purchased from Sigma Mission collection (Basp1: TRCN0000190229 and TRCN0000202255; WT1: TRCN0000054465 and TRCN0000054464). The *Wt1-Kts*overexpression plasmid was generated by the Jaenisch laboratory (Addgene, Plasmid#41082).

The following antibodies were used at a concentration of 1:500 mouse-anti-Oct4 (SantaCruz), rabbit-anti-Nanog (Cosmobio), mouse anti-Sox2 (R&D) and GFP-coupled anti-SSEA-1(ESI-BIO) rabbit anti-Basp1 (Abcam). For *in vitro* differentiation, iPS cells were added onto low attachment plates at 1x10^5^ cells/ml grown in suspension for five days. Embryoid bodies were then plated onto gelatin-coated plates and left to differentiate for approximately two weeks. Differentiated tissues were immunostained using rabbit-α-Tuj1 (Covance), rat-Troma-1 (DSHB), mouse-α-smooth muscle actin (Sigma), m-α-Myosin (Sigma) or m-α-actinin at a concentration of 1:500. To confirm that iPS cell lines harbor the appropriate transgene, PCR was used using primers specific to the sequences in a lentiviral background ^1^(for primer sequences refer to Boland, 2009). All antibodies were validated by testing on positive and negative controls prior to use and/or in the context of an experiment.

### In vitro differentiation

iPSCs were trypsinized and plated (1,000,000 cells/ml) on ultra-low attachment dishes in DMEM supplemented with 10% FBS. To induce neuronal differentiation, EBs were seeded on laminin-coated plates and grown in neuronal differentiation basal media [consisting of DMEM/F-12 (GIBCO)/0.5× N2 (GIBCO)/0.5× B27 (GIBCO)/50 µg/ml BSA fraction V (GIBCO) for an additional 7 days. To induce mesoderm and endoderm differentiation, 4-day EBs (without RA treatment) were seeded on gelatin-coated plates and grown in DMEM (GIBCO) plus 10% FBS (GIBCO) for an additional 7 days. Cells were fixed with 4% paraformaldehyde (Electron microscopy) for 10 min at room temperature. Immunostaining was carried out with standard protocols. The following primary antibodies were used: mouse anti-? III-tubulin (1:500; Covance Research Products) mouse anti-Map2 (Millipore), rabbit anti-GFAP (1:100, Dako);; mouse anti-myosin heavy chain (1:100, Developmental Studies Hybridoma Bank, Iowa City, IA); and goat anti-Sox17 (1:100; <u>R & D Systems</u>, Minneapolis, MN), rat anti-Troma-1 (1:100, Developmental Studies Hybridoma Bank, Iowa City, IA). Antibodies were validated by testing on positive and negative controls prior to use or in the context of the experiment.

### RT-PCR

100,000 cells were plated per 6-well, infected with 0.5 ml/well of the appropriate lentiviruses after 24 hours and transgene expression induced with 5 µg/ml Doxycycline (Sigma) 48 hours after infection. Cells were lysed by adding 1 ml Trizol(tm) Reagent (Ambion) per well and RNA was extracted using the Direct-Zol RNA Mini-Prep kit (Zymogen) according to the manufacturer’s instructions and treated with the provided DNAse1. RNA amounts were normalized with water and converted to cDNA using the iScript(tm) Select cDNA Synthesis Kit (BioRad) using random hexamers or Superscript III (Life technologies) for gene specific priming. 10-20 ng cDNA were amplified in a qPCR reaction using 2x SYBR Select Mastermix (life technologies).

### RNA sequencing

1 µg of RNA was prepped for Sequencing using NEBNext® Ultra(tm) DNA Library Prep Kit for Illumina®. 75 base pair single end reads generated using Illumina’s NextSeq platform were mapped to the mouse genome (mm10) by first removing adapters and low quality bases using Trimmomatic (v0.32, ILLUMINACLIP: TruSeq3-SE.fa:2:30:10 LEADING:3 TRAILING:3).^51^ Reads were then aligned using STAR ^52^ and counts were generated using HTSeq.^53^ RNA-Seq data was analyzed using R an open source programming language and environment for statistical computing and visualization. Differential gene expression analysis was conducted using DESeq2 and gene expression was normalized using vsd transformation to generate heatmaps.^54^

### Immunoprecipitation and Western Blotting

Cells were harvested in complete lysis buffer (20 mM Tris-HCl, 137 mM NaCl, 1% NP-40, 2mM EDTA, protease inhibitor cocktail (Roche)). Protein-G coupled Dynabeads (Novex) were equilibrated with complete lysis buffer for 10 minutes at room temperature before blocking in 5% bovine serum albumin (Fisher Bioreagents) for 10 minutes. Beads were incubated with sample for 2 hour at 4°C, then washed in lysis buffer four times for 10 minutes. Precipitated proteins were eluted from the beads and denatured in Laemmli buffer boiled at 70°C for 10 minutes. Proteins were resolved by SDS-PAGE. Gels were either stained using Coomassie brilliant blue (BioRad) or subsequently transferred onto PVDF membranes (BioRad). Membranes were immunoblotted with anti-Basp1 (Abcam, ab101855), anti-Wt1 (Novus, NBP1-68985) or SoxAb2 and detected with antibody to rabbit IgG (1:1000, A10040-DR546) or antibody to human IgG (1:2000, A11013-DR488). For p-Smad2/3 westerns, MEF cells treated with TGF-β (Cell Signaling) and soluble SoxAb2 or 10 µM RepSox, 10 µM SB431542, 100 ng/ml of TGF-β neutralizing (AB-100NA, R&D systems). Cells were lysed in RIPA buffer (Thermofisher) and incubated on ice with intermittent vortexing for 30 minutes. After the samples were centrifuged at 13,000 X g for 10 minutes, the lysate protein was boiled in LDS sample buffer (Novex) with reducing agent (Novex) at 70°C for 10 minutes, then immediately placed on ice. The samples were loaded in Bolt 4-12% Bis-Tris precast gels (Invitrogen) and later transferred to low fluorescence PVDF membrane (Thermo Scientific). The membranes were blocked with 5% milk in TBST for 2 hours and were immunoblotted with p-Smad 2/3 (1:1000, Cell Signaling #8828) and detected with anti-rabbit IgG HRP (1:3000, Cell Signaling, #7074). Chromatin immunopercipitations were performed using SimpleChip Plus Enzymatic Chromatin IP (Cell signaling) starting with 2 µg of input DNA.

### Validation of shRNA knockdown efficiency by Western blot

MEF cells were transduced with lentivirus and incubated at 37C. Three days after infection the cells were re-suspended and seeded on 6 well plates. On the 4th day 1ug/ml puromycin was added to media, on the 5th day 2ug/ml puromycin was added the media, at 6th day cells were de-attached and counted for viability, MEF cells without lentivirus transduction completely died off after the treatment. Each well of cells were lysed with 50ul lysis buffer, lysate equals to 0.15 million cells were loaded to each lane on protein gel. Membranes were blocked for 1 hour w/ 3% milk at RT, then incubated for two days with primary antibody in 3% BSA at 4C. Antibodies were purchased from Abcam (Anti-BASP1 antibody: ab103315, lot GR315116-4, 1:500 dilution; Anti-Wilms Tumor Protein antibody: ab89901, lot GR177328-39, 1:1000 dilution; Ms mAb to beta Actin [AC-15] (HRP) ab49900; 1:2000 dilution). Membranes were incubated at RT for 30min and washed 3 times, 5min each with PBST before adding the secondary antibody in 3% BSA. (**Fig. S6**)

## Author Contributions

Experiments were initially conceived by R.A.L. and K.K.B. Subsequent experimental design was contributed by all authors and experiments were designed, performed and analyzed by J.W.B. and J.X. Experiments were also performed by S.L., S.G. and N.E. The manuscript was written and revised by J.W.B,J.X. and K.K.B.

### Acknowledgements

We would like to thank J. Gottesfeld, A. Patapoutian, V. Lo Sardo, J. Hazen for assistance and/or helpful discussions, W. Ferguson for technical assistance and K. Spencer for assistance with microscopy. This research was supported by the National Institute on Deafness and other Communication Disorders (DC012592 to K.K.B.), the National Institute of Mental Health (MH102698 to K.K.B.), the California Institute for Regenerative Medicine (RB3-02186 to K.K.B.), the Baxter Family, Norris and Del Webb Foundations (K.K.B.), by Las Patronas and the Dorris Neuroscience Center (K.K.B.), a pre-doctoral fellowship from the California Institute of Regenerative Medicine (J.W.B) the Andrea Elizabeth Vogt Memorial Award (J.W.B.).

